# Deep learning models for identification of splice junctions across species

**DOI:** 10.1101/2021.06.13.448260

**Authors:** Aparajita Dutta, Kusum Kumari Singh, Ashish Anand

## Abstract

Deep learning models like convolutional neural networks (CNN) and recurrent neural networks (RNN) have been frequently used to identify splice sites from genome sequences. Most of the deep learning applications identify splice sites from a single species. Furthermore, the models generally identify and interpret only the canonical splice sites. However, a model capable of identifying both canonical and non-canonical splice sites from multiple species with comparable accuracy is more generalizable and robust. We choose some state-of-the-art CNN and RNN models and compare their performances in identifying novel canonical and non-canonical splice sites in homo sapiens, mus musculus, and drosophila melanogaster.

The RNN-based model named SpliceViNCI outperforms its counterparts in identifying splice sites from multiple species as well as on unseen species. SpliceViNCI maintains its performance when trained with imbalanced data making it more robust. We observe that all the models perform better when trained with more than one species. SpliceViNCI outperforms the counterparts when trained with such an augmented dataset. We further extract and compare the features learned by SpliceViNCI when trained with single and multiple species. We validate the extracted features with knowledge from the literature.

## 1. Introduction

The availability of a plethora of sequenced genomic data has led to the need for structural and functional annotation of the genome. Identification of *splice sites* or *splice junctions* is a crucial step for genome annotation. Splice sites are present at the boundaries of alternating genomic regions called exons and introns where *splicing* occurs [1]. During splicing, the exons and introns in the pre-mRNA clip and ligate in different combinations to form different mature mRNA and eventually form various proteins during translation. The exon-intron junction is called the *donor site* and the intron-exon junction is called the *acceptor site*. Usually, the donor and acceptor sites are characterized by the consensus GT and AG, respectively. Such junctions are called canonical splice junctions. The splice sites that lack the consensus dimers are called non-canonical splice junctions [2].

Machine learning models like support vector machine (SVM), random forest (RF), decision trees (DT), and naïve Bayes (NB) have been applied in the task of splice junction identification [3, 4, 5, 6, 7]. Such models characterize splice junctions in the form of surrounding mono/di/trinucleotide distribution and other positional and density information. However, such manually engineered feature sets are not optimal or exhaustive. This leads to deploying models that can extract relevant features from the genome sequences de novo. With this intention, several deep learning models [8, 9, 10, 11, 12, 13, 14, 15, 16, 17, 18] have been applied to this task in the recent times. Such models can learn and capture the splicing features from the genome sequences by itself.

Among the deep learning models applied, convolutional neural networks (CNN) [8, 9, 11, 16, 17] and recurrent neural networks (RNN) [10, 12, 18] have been applied most frequently to the prediction of splice sites. However, most of the CNN based models and all the RNN based models identify splice sites in a single species. Some of the CNN based models study multiple species like homo sapiens, arabidopsis thaliana, oryza sativa japonica, drosophila melanogaster, and caenorhabditis elegans [9, 16, 17]. However, most of the existing studies do not test the generalizability of the models by training and testing them on novel splice junctions from different species. Furthermore, several existing studies do not contribute towards extraction and interpretation of the non-canonical splicing features learned by the models.

Zuallaert et al. [9] do not discuss the generalization capability of their proposed model by training and testing it on different species. Albaradeia et al. proposed a model named Splice2Deep [17] which trains and tests on different species and still identifies the splice sites with high accuracy. However, they do not extract or discuss any biological features learned by the model. Wang et al. [16] proposed SpliceFinder, which trains a CNN model to identify acceptor, donor, and false splice sites. Although SpliceFinder trains and tests the model on both canonical and non-canonical splice sites of various species, it extracts and visualizes features from canonical sites only. However, a deeper understanding of the non-canonical splicing is equally important as non-canonical splice sites are sometimes vital in regulating important biological events like immunoglobulin gene expression [19]. Furthermore, these CNN based models do not focus on identification of novel splice sites. The ability to identify novel splice sites indicates that a model is generalizable and robust.

To alleviate the limitations of the current research works discussed above, we apply neural network models to identify novel canonical and non-canonical splice sites from various species. In particular, we ask the following questions in this work:

**Question 1:** Whether various neural models perform equally well on identifying splice sites from multiple species? Does any particular model outperform the rest?
**Question 2:** Can the neural models be used to annotate a poorly studied species using data from an extensively annotated species?
**Question 3:** What are the splicing features extracted by the best performing model from the different species?

To answer the above questions, we consider the state-of-the-art RNN and CNN models that have already been applied to identify splice sites. The models are assessed in both the conditions when the training and the testing dataset may be from the same species or different species. We compare the performance of the models to evaluate whether any particular model performs better than the rest in this task. We also augment the dataset by combining training data from more than one species to observe the improvement of the model’s performance in the task of canonical and non-canonical splice site identification. The improvement in a model’s performance on augmenting training data from another species suggests that the model can identify splice sites from poorly annotated species using data from another extensively annotated species. Moreover, all the models are tested on novel splice sites such that the training and testing samples are extracted from two different versions of the dataset.

The state-of-the-art RNN model, called SpliceViNCI, uses a BLSTM network to identify and visualize splice sites within a single species [18]. However, the generalizability of a BLSTM model has not been exploited so far in identifying splice junctions from multiple species. On the other hand, the state-of-the-art CNN models, named SpliceRover and SpliceFinder, have been applied to identify and visualize splice sites from single species [9] as well as across different species [16]. We consider these three neural network models and compare their performances in identifying splice sites across various species.

The contributions of this research can be summarized as:

- We compare CNN and RNN based state-of-the-art models in identifying splice sites from homo sapiens (human), mus musculus (mouse), and drosophila melanogaster (drosophila).
- We evaluate the performance of the models in identifying canonical and non-canonical splice junctions from species on which the models are not trained.
- We evaluate the performance of the models on imbalanced data. Superior performance on imbalanced data indicates that a model is more robust and applicable in the annotation of multiple species.
- We observe that augmenting the training dataset of one species with that of another species improves the performance of all the models, especially in identifying non-canonical splice sites. This indicates that the models can be used to annotate poorly studied species using data from extensively annotated species.
- We extract the splicing features learned from different species by the best performing model and validate them with the existing literature.

## 2. Methods

This section discusses the input representation, network architectures and hyperparameters of the various models used in the study. The visualization technique used for extraction of the biologically relevant features is also discussed. Figure 1 depicts a graphical representation of the workflow common to all the models implemented and the application of visualization.

**Figure 1:**
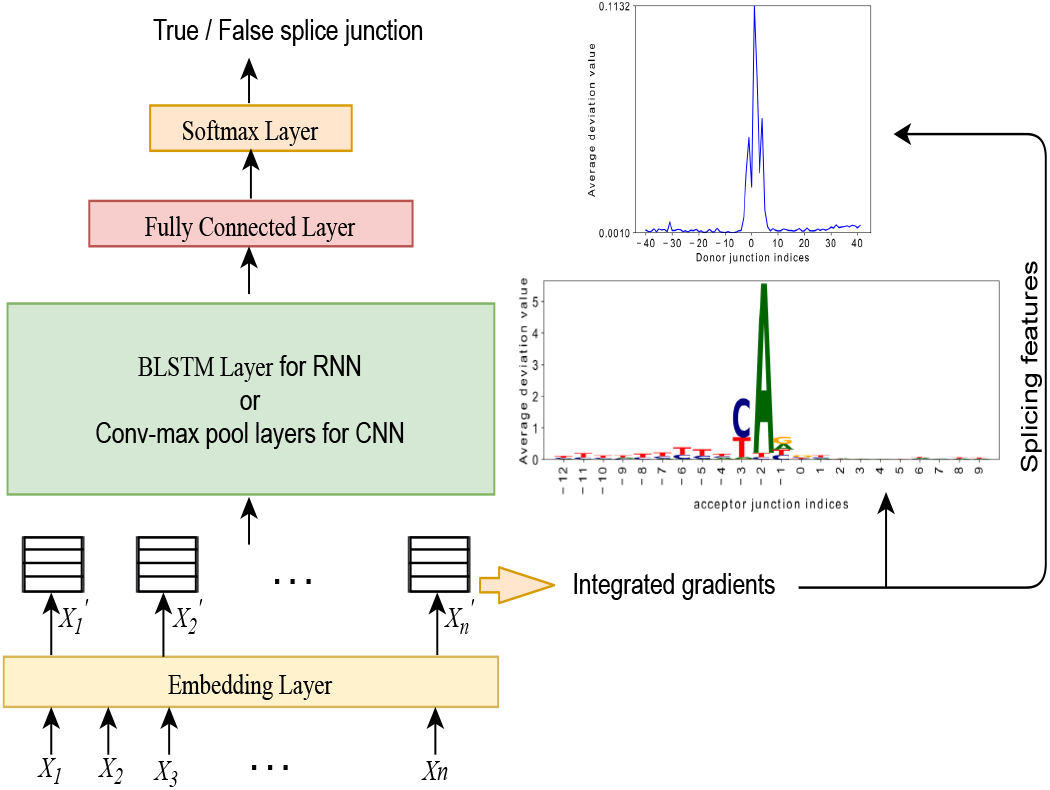
Graphical representation of the workflow.

### 2.1. Input representation

We extract all introns from the protein-coding genes of a given species. The donor and acceptor junction pair corresponding to an intron is an input to the models. Each splice junction is extracted with a flanking upstream and downstream region of 40 nucleotides (nt). The donor and acceptor site sequences corresponding to an intron are concatenated before feeding into the model. These sequences comprise the four nucleotides: *A* (*Adenine*), *C* (*Cytosine*), *G* (*Guanine*), *T* (*Thymine*) and *N* (*denoting any one of the four nucleotides*). Each input sequence comprises 40 nt upstream and downstream regions at both donor and acceptor sites along with the junction dimers. The donor and acceptor site sequences are concatenated to form an input sequence of length 164 nt. This input is fed into each of the model described below.

### 2.2. Models

The following subsections describes the model architectures along with the hyperparameters used for training and testing the models. We partition the training dataset into 90% train and 10% validation data for tuning the hyperparameters.

#### 2.2.1. SpliceRover

This is a CNN model comprising several convolutional layers followed by max-pooling, fully connected, and softmax layers. This model is proposed by Zuallaert et al.[9]. SpliceRover identifies the donor and acceptor splice sites in homo sapiens and arabidopsis thaliana. Additionally, Zuallaert et al. extract the biologically relevant features learned by the model using the DeepLIFT [20] visualization technique.

Zuallaert et al. trained SpliceRover with different numbers of convolutional layers for various datasets of variable sequence length. We trained the model with two convolutional layers followed by a max-pooling, fully connected layer, and softmax layer based on the optimal performance on our dataset. Stochastic gradient descent is chosen as the optimizer with learning rate, decay rate, Nesterov momentum, and the number of steps per learning rate decay set to 0.05, 0.5, 0.9, and 5, respectively. Other hyperparameters like epochs and batch size are set to 30 and 64, respectively.

#### 2.2.2. SpliceFinder

This is a CNN-based model proposed by Wang et al. [16]. The model is trained with the human dataset and identifies donor and acceptor splice sites in several species, namely drosophila melanogaster, mus musculus, rattus, and danio rerio, without retraining. They also extract the relevant splicing features using the DeepLIFT visualization technique.

The model comprises one convolutional layer with 50 kernels of length nine followed by a fully connected layer of size 100. A dropout layer is subsequently added with a dropout rate of 0.3. Finally, there is a softmax layer of 2 nodes to classify true and false splice sites. Adam is used as an optimizer with a learning rate of 10^−4^. Other hyperparameters like epochs and batch size are both set to 50.

#### 2.2.3. SpliceViNCI

This is an RNN-based model comprising an embedding layer, BLSTM layer, fully connected layer, and softmax layer. The model architecture is proposed by Dutta et al. [18]. SpliceViNCI identifies splice junction pairs from the human dataset and visualizes the splicing features by applying visualization techniques like occlusion [21] and integrated gradients [22].

The embedding layer converts each nucleotide into a 4-dimensional dense vector. The subsequent BLSTM, fully connected and softmax layer comprises 100, 1024, and 2 units, respectively. They use binary cross-entropy and Adam [23] as the loss function and the optimizer, respectively. Other tuned hyperparameters are epochs, batch size, dropout, and recurrent dropout, set to 10, 128, 0.5, and 0.2, respectively.

### 2.3. Feature interpretation

We apply Integrated gradients [22] for extracting and interpreting the features learned by the neural model. Integrated gradients is a back-propagation based visualization technique proposed by Sundararajan et al. This technique is based on the computation of gradients along a straightline path from a baseline to the input. The baseline can be a zero embedding vector for text input.

The integrated gradient of an input sequence is given by cumulating the gradients along the straightline path. This cumulative gradient value at each sequence position signifies the importance of that position in the predicted decision of an input. We call the cumulative gradient the deviation value. We consider 50 gradients along the linear path from baseline to input.

The strength of this visualization technique lies in its ability to satisfy two desirable properties of attribution methods: sensitivity and implementation invariance. Intuitively, lack of sensitivity may lead to the attribution method focusing on irrelevant features. Likewise, if an attribution method is not implementation invariant, it may be sensitive to the unimportant biases of the implementation model [22]. Other visualization techniques like DeepLift, Layer-wise relevance propagation (LRP) [24], and gradients lack either of these properties.

## 3. Data

We consider three species, namely *homo sapiens* (*human*), *mus musculus* (*mouse*), and *drosophila melanogaster* (*drosophila*). We choose mouse and drosophila species in order to have a wide range of comparison because mouse is closer to human in the phylogenetic tree, whereas drosophila is further [25]. Mouse and human species have 99% homologous protein-coding genes [26]. In contrast, drosophila and human have 60% homologous protein-coding genes [27]. The procedures of generating the positive and negative data are described in the next section.

### 3.1. Positive data

The genome sequence data (FASTA files) and the corresponding annotations (GTF files) are downloaded from the GENCODE database [28] for human and mouse species. The FASTA and GTF files for Drosophila melanogaster were downloaded from the Ensembl database [29].

We test the various models on the identification of novel splice sites. Therefore, the training and testing samples are extracted from two different versions of the database. The training data is extracted from an earlier released version. The testing data is generated from a later release such that the testing samples are not present in the training version of the database. This ensures that the model can identify splice sites that were not annotated in the training version of the dataset, thus making the model more robust.

The number of introns present in the training and testing data of the different species is mentioned in Table 1 along with their corresponding reference genome and release versions. We observe that the number of introns in drosophila is much less than in human and mouse. This is because the drosophila genome comprises only four pairs of chromosomes compared to 23 and 20 pairs in human and mouse species, respectively.

**Table 1:** Distribution of positive dataset in mouse, human and drosophila.

| Species (reference genome) | Training data | Testing data |
| --- | --- | --- |
|  | # of introns (version) | # of introns (version) |
| *Mus musculus* (GRCm38) | 225616 (V:M2) | 26030 (V:M24) |
| *Homo sapiens* (GRCh38) | 290502 (V:20) | 5612 (V:26) |
| *Drosophila melanogaster* (BDGP6) | 58522 (V:95) | 118 (V:103) |

### 3.2. Negative data

The negative dataset is generated by randomly extracting genome sequences from the genome sequence data. We randomly search for the donor site dimer and a subsequent acceptor site dimer. Both the donor and acceptor site dimers are present in the same chromosome and are not annotated as splice sites in the positive dataset. The donor-acceptor dimer pairs considered in the generation of negative dataset are *GT-AG, GC-AG*, and *AT-AC*.

The *GT – AG* consensus rule applies to all the three species considered here [30, 31]. *GC – AG* and *AT – AC* are the most frequently occurring non-canonical dimer pairs in mouse species of our positive data. This corroborates the most frequently occurring non-canonical splice junctions in the human species known from literature [32]. The most frequent non-canonical dimer pair in our drosophila positive data is *GC – AG* and *GT – TG*, closely followed by *AT – AC*. However, for the uniformity of comparison, we use *GC – AG* and *AT – AC* as non-canonical negative dimer pairs across all species. The non-canonical data comprises 1% *GC – AG* and 1% *AT – AC* splice junctions. This proportion is chosen based on a study by Stephen M Mount [32]. We extract the splice sites along with a flanking upstream and downstream region equal to that of the positive samples.

A major challenge in identifying splice sites lies in the fact that although GT-AG is the most frequently occurring splice junction dimer pair, there are also instances of splice sites that lack this dimer pair at the junctions. This suggests the presence of additional splicing signals in the vicinity of the sites which govern the splicing phenomenon. We add the non-canonical dimer pairs in the negative dataset to make the model capable of recognizing such splicing signals present near the non-canonical splice junctions, which differentiate the true and decoy splice sites.

## 4. Results

The results obtained from various analysis and comparison of the models are discussed in the following subsections.

### 4.1. SpliceViNCI outperforms other models in identifying splice sites of multiple species

We intend to evaluate the performance of various neural network models on the task of identifying canonical and non-canonical splice sites from multiple species. We use accuracy, precision, recall, and F1-scores as the evaluation metrics. We train and test SpliceViNCI, SpliceRover and SpliceFinder prediction models with data from human, mouse and drosophila.

Figure 2 displays the performances of SpliceViNCI, SpliceFinder, and SpliceRover in the identification of canonical and non-canonical splice sites where each model is trained and tested with the data from the same species. We observe that SpliceViNCI outperforms both the CNN models in most of the metrics for all three species. This analysis is done to test the consistency of the models in the identification of splice sites in various species.

**Figure 2:**
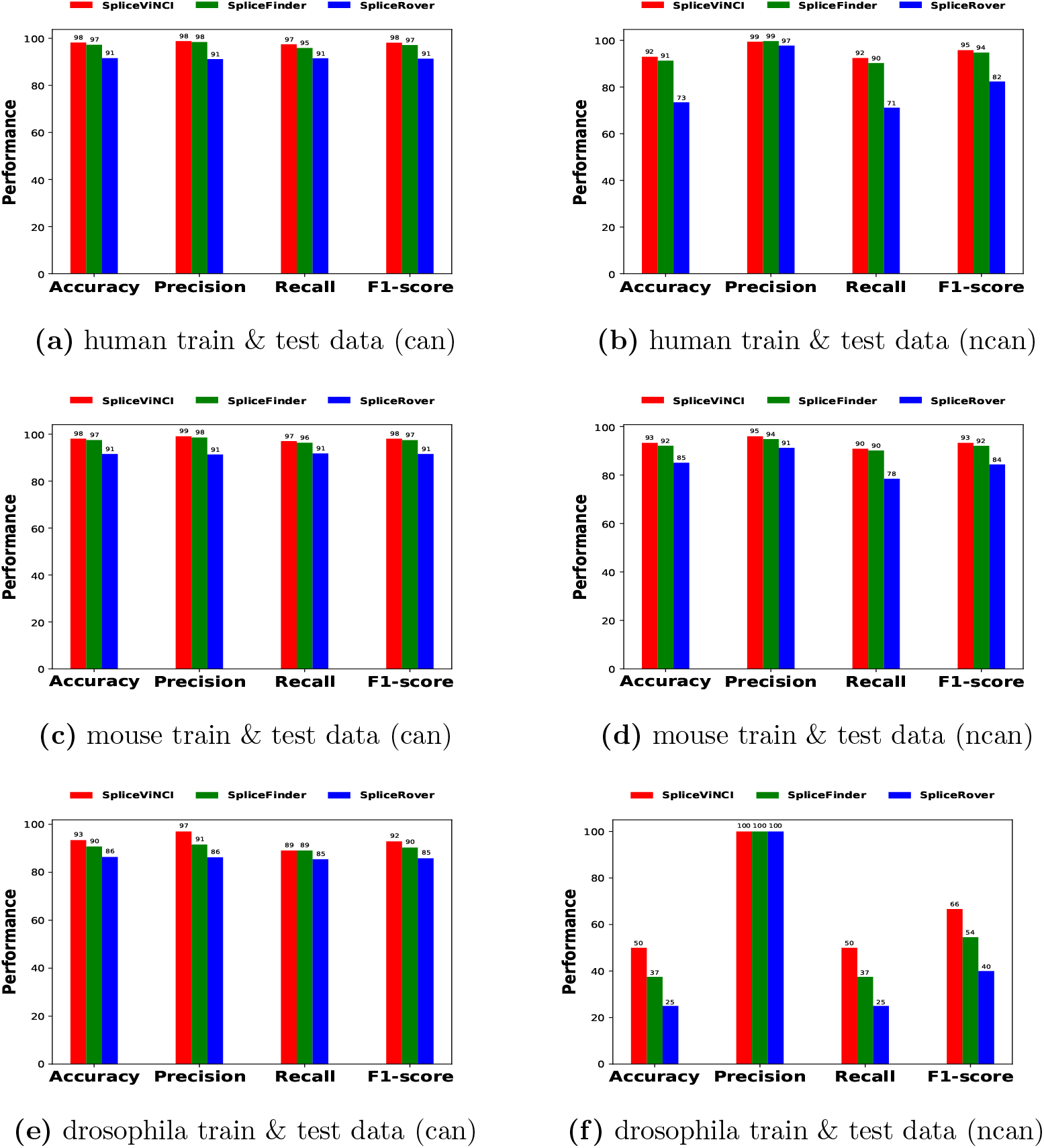
Performance (in percentage) obtained by SpliceViNCI, SpliceRover and SpliceFinder in the identification of canonical (can) and non-canonical (ncan) splice junctions when trained and tested with data from the same species.

We observe in Figure 2 that SpliceViNCI obtains an F1-score approximately 1% to 2.5% more than SpliceFinder in the identification of canonical splice junctions across the three different species. Furthermore, The F1-score of SpliceViNCI is 1% to 12% more than SpliceFinder in identifying non-canonical splice sites. The F1-score of SpliceViNCI is 7% (13% to 26%) more than SpliceRover in the identification of canonical (non-canonical) splice junctions. Similarly, SpliceViNCI outperforms SpliceFinder and SpliceRover in the other three performance metrics: accuracy, precision, and recall.

Additionally, we observe a relatively poor performance of all the three models in identification of non-canonical splice sites in drosophila. This can be attributed to the selection of *GC – AG* and *AT – AC* as non-canonical negative dimer pairs in the training dataset. Whereas, in the real scenario, the most frequent non-canonical dimer pair in our drosophila positive data is *GC – AG* and *GT – TG*. This difference of non-canonical dimer pairs in the positive and negative data of drosophila may lead to the model missing out the subtle splicing signals present in the vicinity of the most frequently occurring non-canonical dimer pairs.

### 4.2. SpliceViNCI outperforms other models in identifying splice sites of unseen species

Next we were curious to test the generalizability of the models in recognizing splice sites across different species. For this objective, we trained and validated each model with only human data and tested them on mouse and drosophila. In other words, this analysis tests the performance of the models on unseen species.

Figure 3 shows the performance of SpliceViNCI, SpliceFinder, and SpliceRover trained with human data in identifying canonical and non-canonical splice junctions from mouse and drosophila test data. SpliceViNCI obtains an F1-score up to 4% and 12% more than SpliceFinder in the identification of canonical (Figure 3(a)) and non-canonical (Figure 3(b)) splice junctions, respectively. On the other hand, F1-score obtained by SpliceViNCI is up to 10% and 12% more than SpliceRover in the identification of canonical (Figure 3(c)) and non-canonical (Figure 3(d)) splice junctions, respectively. This test further affirms the robustness of SpliceViNCI compared to the counterparts.

**Figure 3:**
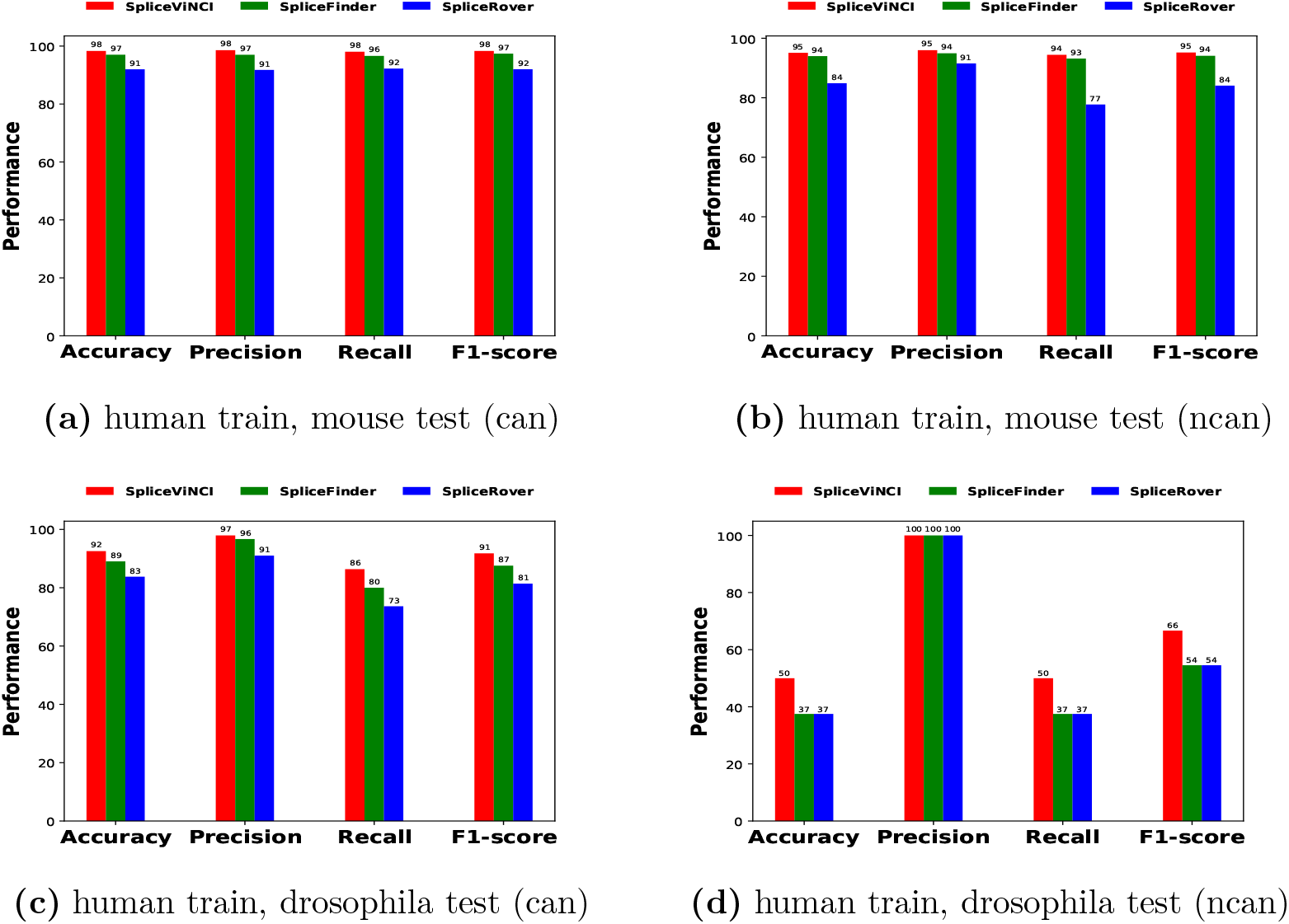
Performance (in percentage) obtained by SpliceViNCI, SpliceRover and SpliceFinder trained with human data in the identification of canonical (can) and non-canonical (ncan) splice junctions from mouse and drosophila test data.

### 4.3. SpliceViNCI is more robust with imbalanced training data

We test the robustness of the neural network models by training the models on an imbalanced dataset. An imbalanced dataset is generated by increasing the ratio of negative to positive samples. This ratio of negative to positive samples is called the decoy rate. We increase the decoy rate from 3 to 9 in an interval of 2. Since accuracy, precision and recall are not appropriate performance metrics in an imbalanced dataset; we display only F1-score in this analysis.

We train the models on the human dataset and test them on the human, mouse, and drosophila datasets. The positive and negative training samples are extracted from chromosome 1 (chr1) of the human dataset. We chose chromosome 1 since it is the largest chromosome in the human genome. Figure 4 displays the F1-scores obtained by SpliceViNCI, SpliceFinder, and SpliceRover when trained on an imbalanced human dataset and tested on canonical and non-canonical splice sites from different species.

**Figure 4:**
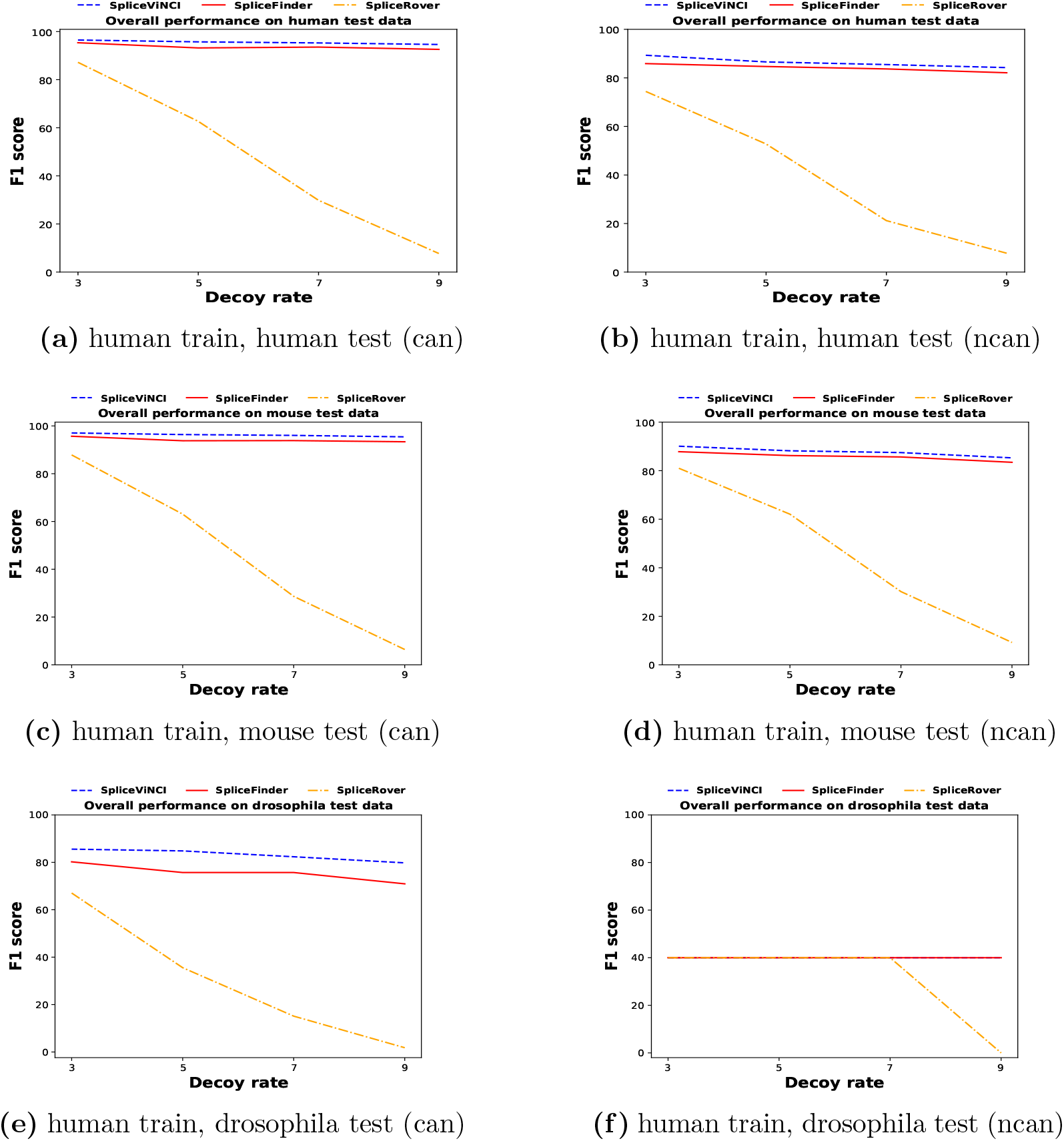
F1-score (in percentage) obtained by SpliceViNCI, SpliceFinder and SpliceRover in the identification of splice junctions with imbalanced training data.

We observe that both SpliceViNCI and SpliceFinder maintain its performance with increasing decoy rate whereas the performance of SpliceRover significantly reduces by 80% as the decoy rate increases from 3 to 9. However, SpliceViNCI consistently outperforms SpliceFinder and SpliceRover across different species even when the decoy rate increases. In particular, the performance of SpliceRover reduces to less than 50% when the decoy rate increases beyond 5. On the contrary, the performance of SpliceFinder and SpliceViNCI reduces up to 10% and 6%, respectively. Therefore, we can conclude that SpliceViNCI is the most robust model across several species to identify canonical and non-canonical splice sites.

### 4.4. SpliceViNCI outperforms other models in identifying splice sites of partially annotated species

We investigate whether the performance of the models can improve on augmenting the training data from one species with training data from another. If a model performs better with such an augmented dataset, it can then be applicable to annotate partially studied species using data from extensively annotated species. With this purpose, we extract training samples from chromosome 1 for the human and mouse training data. The training samples for drosophila were extracted from chromosome 3R. We choose these chromosomes since they are the largest chromosomes in their respective genome sequence data.

Figure 5(a) (Figure 5(b)) shows the performance of SpliceViNCI, SpliceFinder, and SpliceRover in the identification of canonical and non-canonical splice sites when trained with only mouse (drosophila) data and mouse (drosophila) data combined with human data. We observe that all three models display improvement in the F1-score when trained using data from two species. This improvement is observed in the case of both canonical and non-canonical splice site identification.

**Figure 5:**
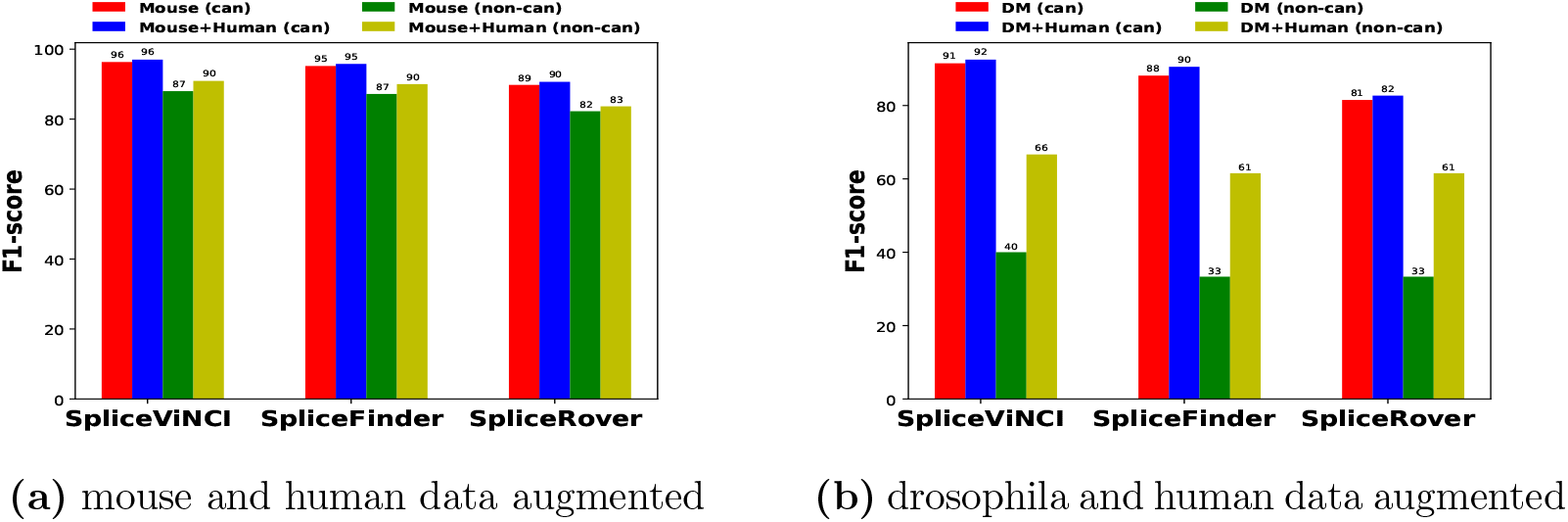
F1-score (in percentage) obtained by SpliceViNCI, SpliceFinder and SpliceRover in the identification of splice junctions with single species and multi-species training data.

Hence, we can infer that combining the training data of one species with samples from another extensively annotated species can improve the model’s performance. This performance improvement can be attributed to the increase in the number and variation of training data when multiple species are used. This analogy can also motivate annotating poorly studied or newly annotated species by training a model with data from another extensively annotated species.

In canonical splice sites, all three models show an improvement of approximately 1% when human data is added for training the model instead of using only mouse or drosophila data. However, in non-canonical splice sites, the improvement is up to 28% when human data is added to the training dataset. The more significant improvement in identifying non-canonical splice sites can be attributed to the fact that canonical splice site motifs are primarily similar across all eukaryotes [16, 30]. On the other hand, non-canonical splice sites show a wider range of variations and frequencies of occurrence. Therefore higher variation in the training data assists the models to recognize more unseen non-canonical splice sites.

Furthermore, we observe that SpliceViNCI performs better than SpliceFinder and SpliceRover in both the scenario when one or two species are used to train the model. SpliceViNCI shows an improvement of up to 7% compared to SpliceFinder when only mouse or drosophila data is used for training the models. The performance of SpliceViNCI improves up to 5% compared to SpliceFinder when human data is also used for training. The improvement of SpliceViNCI compared to SpliceRover goes up to 10%.

### 4.5. SpliceViNCI captures significant sequence positions

As observed in Figure 5, combining data from two species for training the models significantly improves the performance in non-canonical splice sites. Furthermore, SpliceViNCI shows the highest improvement when such an augmented data is used. Therefore, we extract the features captured by SpliceViNCI from the non-canonical mouse and drosophila test data with and without augmentation and compare the findings from both. We represent the augmented dataset from two different species *S*_1_ and *S*_2_ in the form of *S*_1_ + *S*_2_.

#### 4.5.1. Significant sequence positions captured in mouse

Figure 6(a) and Figure 6(c) display the importance of sequence positions captured for non-canonical donor sites by SpliceViNCI when trained by mouse+human and mouse data, respectively. We observe that the mouse model captures only the splice site and its downstream region as significant. In contrast, the mouse+human model captures the upstream region of the donor site as significant as well. Since the mouse and human splice site consensus at donor and acceptor sites are conserved [30, 31], we can derive that the mouse+human model captures the extended upstream region of the donor site consensus 9-mer [*AC*]*AGGTRAGT* whereas the mouse model does not.

**Figure 6:**
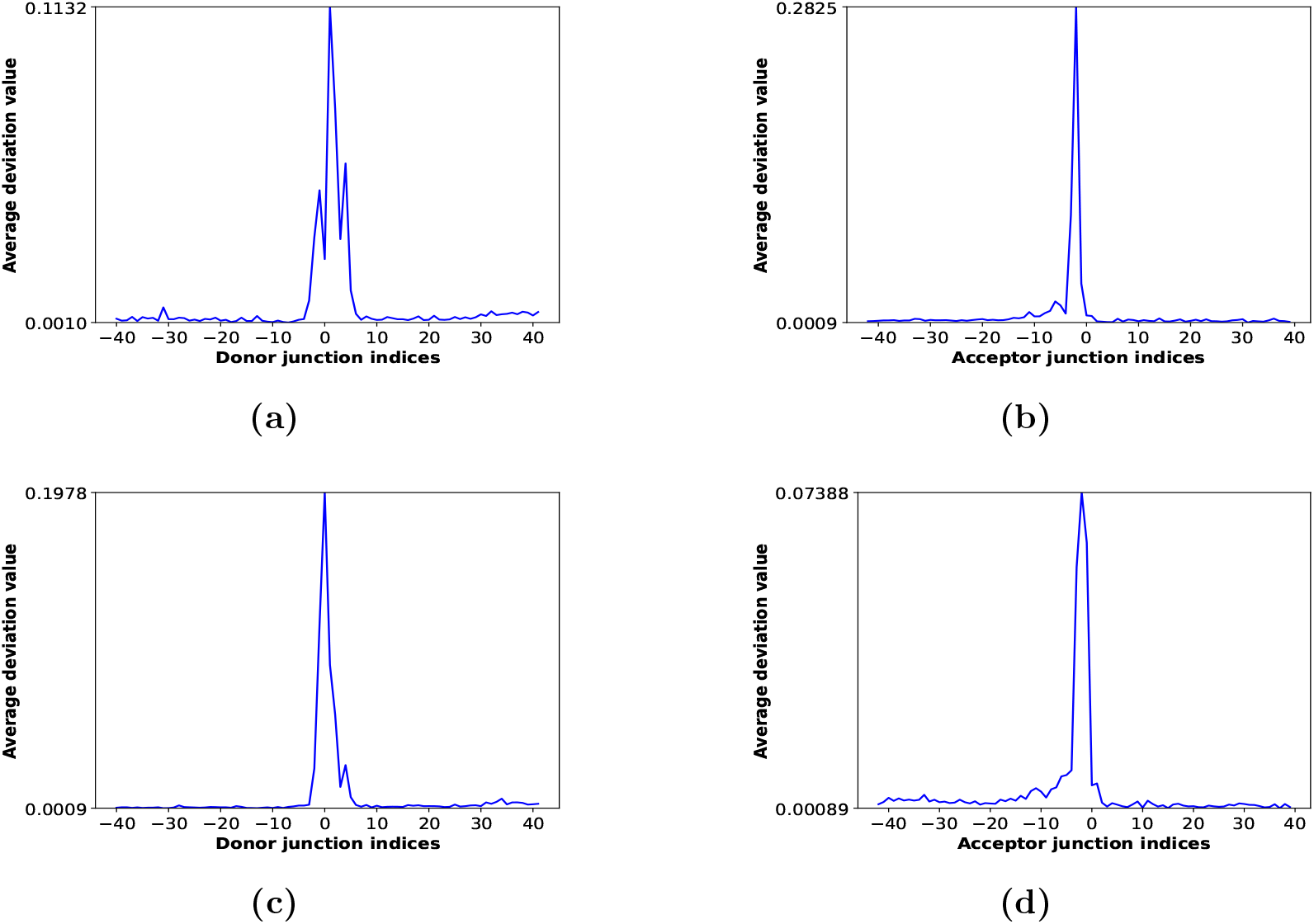
The importance of sequence positions. The average deviation value per position obtained by integrated gradients for non-canonical (a) donor junction and (b) acceptor junction in mouse+human model; non-canonical (c) donor junction and (d) acceptor junction in mouse model.

The significant positions captured by both the models for the acceptor junction (Figure 6(b) and Figure 6(d)) seem to be identical. However, the importance of sequence positions beyond −15 nt upstream of the acceptor site appears smoother when mouse+human data (Figure 6(b)) is used. Beyond −15 nt upstream of the acceptor junction is the position where the polypyrimidine tract ends.

#### 4.5.2. Significant sequence positions captured in drosophila

The importance of sequence positions at the donor sites of drosophila is depicted in Figure 7(a) and Figure 7(c) for drosophila+human and drosophila model, respectively. We observe that the drosophila+human model gives maximum importance to position [0] at the donor junction compared to the extended upstream and downstream region. On the contrary, the drosophila model gives more importance to the upstream region than the donor junction.

**Figure 7:**
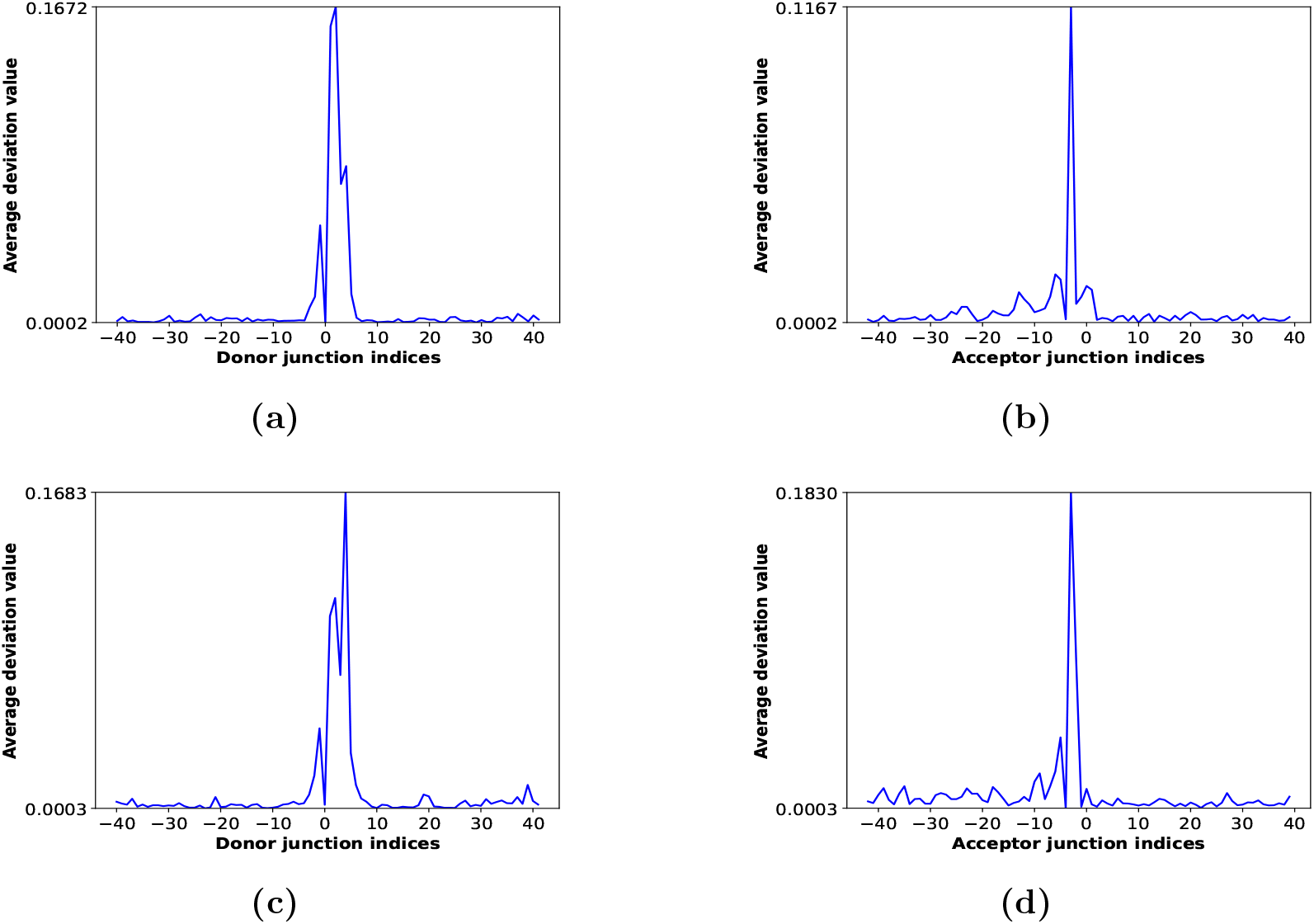
The importance of sequence positions. The average deviation value per position obtained by integrated gradients for non-canonical (a) donor junction and (b) acceptor junction in drosophila+human model; non-canonical (c) donor junction and (d) acceptor junction in drosophila model.

As known from the literature, the consensus dimer at the junctions differentiate non-canonical splice sites from their canonical counterparts. Intuitively, the importance given by the drosophila+human model (Figure 7(a)) abides by this rule thus making more sense than the drosophila (Figure 7(c)) model. In the case of acceptor junctions, drosophila+human (Figure 7(b)) model captures downstream region of the junction which the drosophila ((Figure 7(d))) model does not. Furthermore, the region captured beyond the PY-tract appears smoother in the drosophila+human model.

### 4.6. SpliceViNCI captures splice junction consensus

We observe that SpliceViNCI captures donor and acceptor splice site motifs in the case of both mouse+human (Figure 8(a) and Figure 8(a)) and mouse (Figure 8(c) and Figure 8(d)) training data. However, on a closer observation, we see that the donor site motif obtained from mouse+human training data (Figure 8(a)) covers more extended consensus compared to that of the mouse training data (Figure 8(c)).

**Figure 8:**
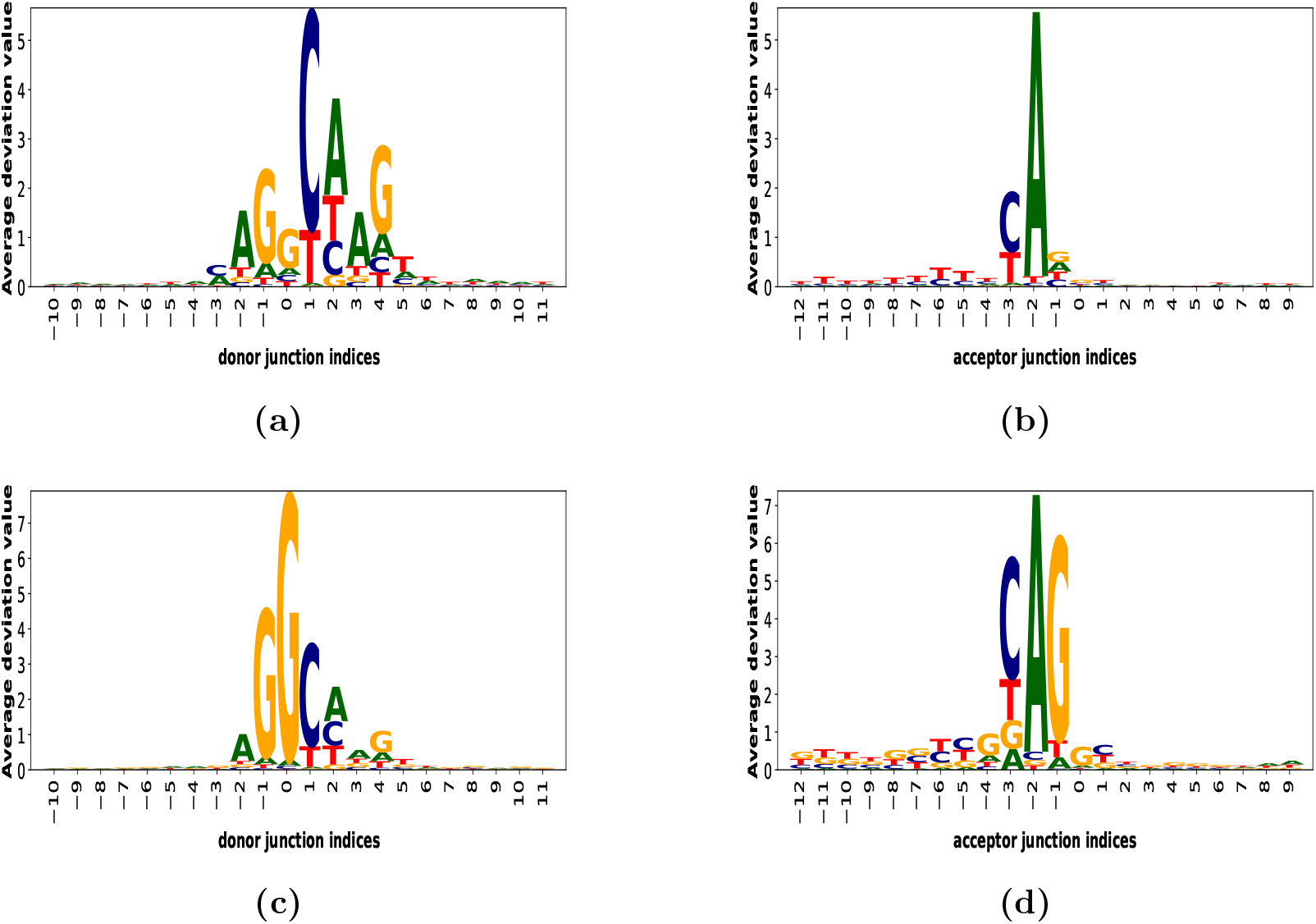
Sequence motifs at splice junctions. The average deviation value per position per nucleotide is shown by integrated gradients for non-canonical (a) donor junctions and (b) acceptor junctions in mouse+human model; non-canonical (c) donor junctions and (d) acceptor junctions in mouse model.

Additionally, The mouse+human model gives highest importance to position [0] which corresponds to the most common non-canonical donor site consensus [GC]. The PY-tract captured by mouse+human model (Figure 8(b)) is less noisy compared to that of the mouse model (Figure 8(d)). Smoothening of the PY-tract is also seen in the drosophila+human model (Figure 9(d)) compared to drosophila model (Figure 9(b)).

**Figure 9:**
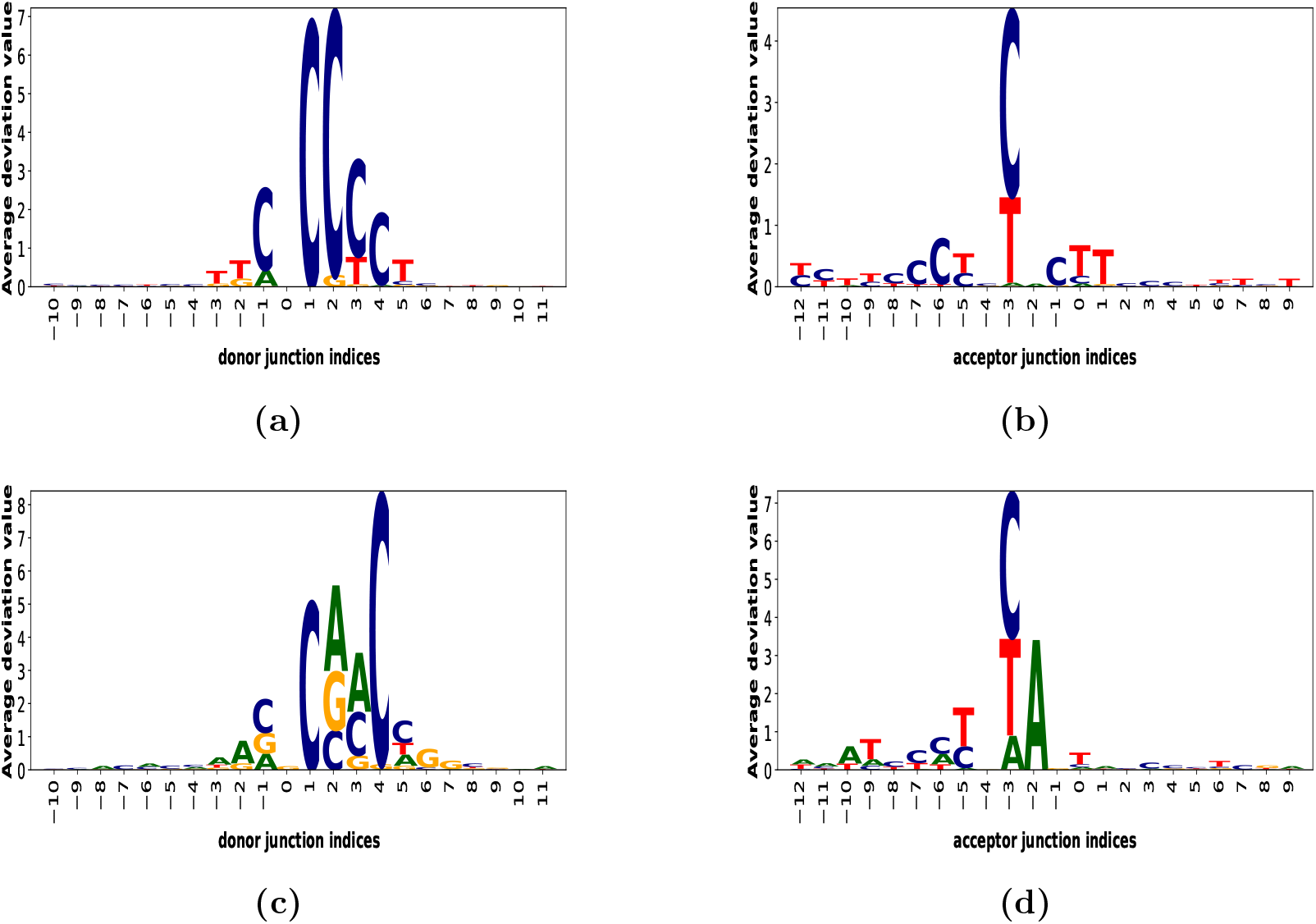
The importance of nucleotides per sequence position. The average deviation value per nucleotide per position is shown by integrated gradients for non-canonical (a) donor junction and (b) acceptor junction in drosophila+human model; non-canonical (c) donor junction and (d) acceptor junction in drosophila model.

## 5. Conclusion

CNN and RNN models have been frequently applied to identify splice sites. However, most of the applications do not evaluate the performance of the models across multiple species or unseen species. We select some state-of-the-art CNN (SpliceRover and SpliceFinder) and RNN models (SpliceViNCI) that have already been applied to identify splice sites. We compare the performances of the models in identifying novel canonical and non-canonical splice sites in human, mouse, and drosophila species.

SpliceViNCI outperforms other state-of-the-art models in identifying the splice junctions in all the three species. SpliceViNCI also outperforms its counterparts in identifying splice sites from species on which it is not trained. Furthermore, SpliceViNCI attains the highest F1-score when trained with an imbalanced dataset. The above analysis suggest SpliceViNCI as a more robust and generalizable model than its counterparts.

We observe an improvement in the performance of all the models when data from multiple species are used for training. SpliceViNCI performs better than the counterparts with such augmented training data as well. Therefore, SpliceViNCI proves to be a preferable choice for annotating species that are either newly or poorly annotated using training data from extensively annotated species.

We further extract the splicing features learnt by SpliceViNCI through the application of integrated gradients. The knowledge thus obtained is validated with the existing literature. We also compare the features extracted when the model is trained with single-species and multiple-species training data. We observe that SpliceViNCI extracts more specific features when trained using data from more than one species.

## Acknowledgments

K.K. Singh acknowledges the grant from SERB (CRG/2019/001352). We acknowledge the Department of Biotechnology, Govt. of India for the financial support for the project BT/COE/34/SP28408/2018.

